# Mind reading of the proteins: Deep-learning to forecast molecular dynamics

**DOI:** 10.1101/2020.07.28.225490

**Authors:** Chitrak Gupta, John Kevin Cava, Daipayan Sarkar, Eric Wilson, John Vant, Steven Murray, Abhishek Singharoy, Shubhra Kanti Karmaker

## Abstract

Molecular dynamics (MD) simulations have emerged to become the backbone of today’s computational biophysics. Simulation tools such as, NAMD, AMBER and GROMACS have accumulated more than 100,000 users. Despite this remarkable success, now also bolstered by compatibility with graphics processor units (GPUs) and exascale computers, even the most scalable simulations cannot access biologically relevant timescales - the number of numerical integration steps necessary for solving differential equations in a million-to-billion-dimensional space is computationally intractable. Recent advancements in Deep Learning has made it such that patterns can be found in high dimensional data. In addition, Deep Learning have also been used for simulating physical dynamics. Here, we utilize LSTMs in order to predict future molecular dynamics from current and previous timesteps, and examine how this physics-guided learning can benefit researchers in computational biophysics. In particular, we test fully connected Feed-forward Neural Networks, Recurrent Neural Networks with LSTM / GRU memory cells with TensorFlow and PyTorch frameworks trained on data from NAMD simulations to predict conformational transitions on two different biological systems. We find that non-equilibrium MD is easier to train and performance improves under the assumption that each atom is independent of all other atoms in the system. Our study represents a case study for high-dimensional data that switches stochastically between fast and slow regimes. Applications of resolving these sets will allow real-world applications in the interpretation of data from Atomic Force Microscopy experiments.

## 1 Introduction

Molecular dynamics or MD simulations have emerged to become the cornerstone of today’s computational biophysics, enabling the description of structure-function relationships at atomistic details[19]. These simulations have brought forth milestone discoveries including resolving the mechanisms of drug-protein interactions, protein synthesis and membrane transport, molecular motors and biological energy transfer, and viral maturation, encompassing a number of our contributions[9]. More recently, we have employed molecular modeling to predict mortality rates from SARS-Cov-2[26], showcasing its application in epidemiology.

In MD simulations, the chronological evolution of an *N*-particle system is computed by solving the Newton’s equations of motion.

Methodological developments in MD has pushed the limits of computable system-sizes to hundreds of millions of interacting particles, and timescales from femtoseconds (10^*−*15^ second) to microseconds (10^*−*6^ second), allowing all-atom simulations of an entire cell organelle [23]. High performance computing, parallelized architecture, speciality hardware and GPU-accelerated simulations have made notable contributions towards this progress. However, in spite of significant advancements in both development and applications, computational resources required to achieve biologically relevant system-sizes and timescales in brute-force MD simulations remain prohibitively “expensive”. Notably, MD involves solving Newtonian dynamics by integrating over millions of coupled linear equations. An universal bottleneck arises from the time span chosen to perform the numerical integration. Akin to any paradigm in dynamic systems, the time span for numerical integration is limited by the dynamics of the fastest mode. In biological systems, this span is 2 femtoseconds (fs) or lower, owing to the physical limitations of capturing fast vibrations of hydrogen atoms. Thus, MD simulation of at least 1 microsecond, wherein biologically relevant events occur, requires the computation of 500 million fs-resolved time steps. Each step involves the calculation of the interaction between every particle with its neighbors, which scales as *N^2^* or *N* log*N*. When *N* = 1-100 million atoms, these simulations are only feasible on peta to exascale supercomputers.

Several techniques have been employed to accelerate atomistic simulations, which can broadly be classified into two categories: coarse-gaining and enhanced sampling. In the former, the description of the system under study is simplified in order to reduce the number of particles required to completely define the system[9]. In the latter, either the potential energy surface and gradients (or forces) that drive the molecular dynamics is made artificially long-range so as to accelerate the movements or multiple short replica of the system are simulated in order to sample a broader range of molecular movements than a long brute-force MD[13]. A major contention of these techniques is that, the simulated protein movements cannot be attributed either chemical precision or a realistic time label[9].

We explore machine-learning methodologies for predicting the outcomes of MD simulations by preserving their accurate time labels. This idea will greatly reduce the computational expenses associated with performing MD, making it broadly accessible beyond the current user-base of scientific researchers to high schools and colleges, where the computational resources are sparse. The developments will imminently expedite the efforts of nearly 20,000 users of our open-source MD engine NAMD[19]. In this resource paper, we present two types of data sets, the dynamic correlations within which pose significant challenge on existing machine-learning techniques for predicting the real-time nonlinear dynamics of proteins. The underlying physics of these data sets represent out-of-equilibrium and in-equilibrium conditions, wherein the *N*-particle systems evolve in the presence *vs.* absence of external perturbations. Beyond tracking the nonlinear transformations, these examples also create an opportunity to study whether prediction accuracy of future outcomes with fs-resolution improve, if prior knowledge is utilized to enhance the signal-to-noise ratio of key features in the training set.

A number of works in the past have focused on predicting protein structures from protein sequence/ compositional information by training on the so-called *sequence-structure relationship* using massive data sets accrued over the PDB and PSI data-bases[2]. However, knowledge of stationary 3D coordinates offer little to no information on how the system evolves in time following the laws of classical or quantum Physics. Little data is available to train algorithms on such time series information despite the imminent need to predict molecular dynamics[15]. The presented data sets capture both the linear and nonlinear movements of molecules, resolved contiguously across millions of time points. These time series data enable the learning of spatio-temporal correlation or memory-effect that underpins Newtonian dynamics of large biomolecules - a physical property that remains obscure to the popular sequence-structure models constructed stationary data. We establish that the success of any deep learning strategy towards predicting the dynamics of a molecule with fs precision is contingent on accurately capturing on these many-body correlations. Thus, the resolution of our MD data sets will result in novel training strategies that decrypt an inhomogeneously evolving time series. As a publicly accessible resource, our MD simulations trajectories of even larger systems (10^5^-10^7^ particles)[23] will be provided in the future to seek generalizable big-data solutions of fundamental Physics problems.

In what follows, we use equilibrium and non-equilibrium MD to create high-dimensional time series data with atom-scale granularity. For simplicity, we derive a sub-space of intermediate size composed only of carbon atoms. In this intermediate-dimensional space, where the data distribution is densed highly correlated, we train state-of-the-art time sequence modeling techniques including recurrent neural networks (RNNs) with long short term memory (LSTMs) to predict the future state of the system (Fig. 1). We explore, how a Kirchhoff decomposition[1] of the many-body problem dramatically enhances the learning accuracy both under equilibrium and non-equilibrium data, even when the number of hidden layers *<<* than the number of atoms. Hardness of the time series are captured in terms of root mean square deviations (RMSD) errors, computed at different lead-times. The RMSD between two *N*-dimensional data points A and B is defined as:

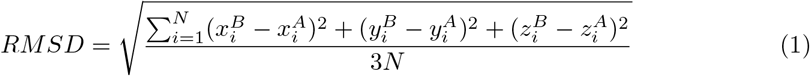

where A and B could be either real and predicted points. We also define *history time* and *lead time* to be a moving window of cumulative time steps (in units of fs) respectively in the *past* and in the *future* of a given data point in the time series, over which training and predictions are achieved. Modeling accuracy was evaluated by varying the amount of historical data points incorporated during the training phase, and then comparing its prediction accuracy against that of a static or linear model. Surprisingly, we find that the equilibrium MD time series is more challenging to learn, despite the non-Gaussian distribution atoms associated to the non-equilibrium MD. Henceforth, we discuss how these new data-set resources can be used for future research of modeling high-dimensional high-frequency event-driven MD time series data.

**Fig. 1:**
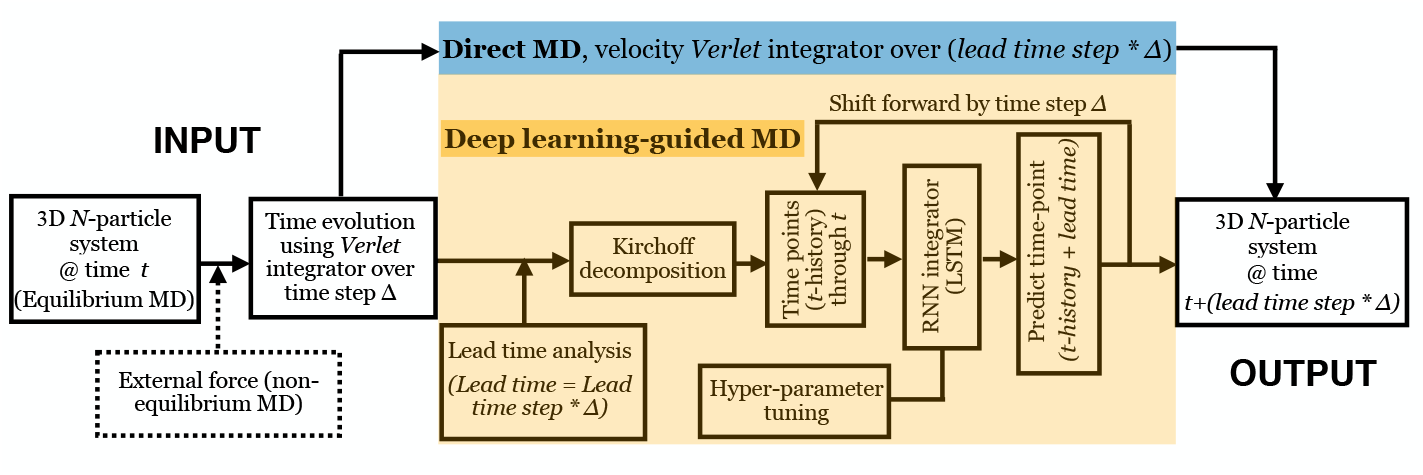
Work-flow of classical and data-guided MD simulations. Classical MD simulations use the Verlet integrator to march forward in time with finite step size ∆ (blue). Deep learning guided MD, uses features from MD, learns using a neural network to forecast future rare events (yellow).

## 2 Related Works

In the recent past, machine learning approaches have been successful in analyzing the results of MD simulations. Support vector machines and variational auto-encoders have been developed to extract free energy information from MD simulations[15]. Kinetic properties of small-molecules have also been extracted using neural networks. It is also shown, that neural networks trained on limited data selected from very expensive MD simulations can resurrect the entire Boltzmann distribution for small proteins[15]. However, none of these approaches are aimed at resurrecting the real-time (i.e. fs-resolution molecular movements of biological molecules) – one of the central goals of MD simulations [19]. RNNs and LSTMs have been used to predict MD [5], but the tested data sets fail to wholly capture the dynamical complexity of a biological molecule. A key observation made therein that inspires our current investigations is that training on molecular dynamics beyond 16 particles is improbable. The data sets we present in the next section challenges this seminal bottleneck that must be overcome to forecast MD simulations of real biological systems.

From a computational perspective, any dynamically evolving system can be regarded as event-driven time series data; in this sense, MD simulations are essentially high-dimensional high-frequency time series data, and sequence modeling techniques like Recurrent Neural Networks [4], Hidden Markov Models and ARMA, can be used to model MD trajectories. Deep learning has recently emerged as a popular paradigm for modeling dynamically evolving time series and predicting future events. These techniques have also been vastly studied in special application areas like business and finance [21], healthcare [14], power and energy [20].

## 3 Problem Formulation

At room temperature, where biology exists, Newtonian mechanics of the molecules become stochastic described by the fluctuation-dissipation theorem. The ensuing molecular trajectories converge at Boltzmann-distributed ensembles at infinitely long times. It has been established that protein dynamics in cells can be modeled as motions of molecules within a media that is highly viscous. Imposing this so called *friction-dominated* condition on the stochastic Newton’s equations, and assuming that a *complete set* of the degrees of freedom for describing the dynamical system is known, molecular dynamics is deemed to be a Markovian process. In simpler terms, it is a process for which predictions can be made regarding future outcomes based solely on its present state, and most importantly, such predictions are just as good as the ones that could be made knowing the process’s full history. The equation of motion of a particle of mass (*m*), at position (*x*) in time (*t*) within an environment of friction coefficient (*γ*) becomes:

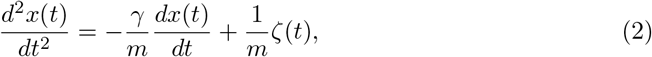

where the random force *ζ* is constrained by requiring the integral of its autocorrelation function to be inversely related to the friction coefficient.

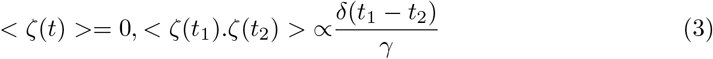

However, we often cannot find a complete set of descriptors to probe the molecular dynamics of proteins. The problem becomes particularly challenging once the number of amino acids in the protein sequences becomes more than 115[17] (i.e. roughly *N* = 1150 atoms). The associated phase space (of 3*N* positions, *X* = *x*_1_*, x*_2_*, &x_N_*, and 3*N* momenta) for systems of these sizes (or higher) becomes too extended for physics-based methods such as MD to visit all the possible points in the 6*N*-dimensional space. This incomplete description of the phase space together with the well-known finite-size artifacts[19], introduces “memory” into any realistic MD simulation. Introduced originally by Zwanzig and used in ref. [22], this memory shows up as a “long-time” tail in auto-correlation functions of atoms undergoing simulation. In a fully equilibrated systems, this memory is short-term vanishing within picoseconds (10^*−*12^ seconds) for carbon, hydrogen and oxygen atoms that primarily compose the proteins[7]. In non-equilibrium simulations that are often employed to accelerate MD[10], the long-time tail stretches to nanosecond (10^*−*9^ seconds). Noting that every integration time step in MD is 1-2 fs (10^*−*15^ seconds), there exists at least 6 order of magnitude in time within which the memory of the system is relevant and offers the opportunity to leverage deep learning techniques for making predictions.

Computational modeling of any complex dynamics essentially boils down to a multivariate time series forecasting task, and hence time series trajectory data capturing an evolving biological system is necessary to analyze and computationally learn the underlying molecular dynamics. Below we first present some basic definitions and notations we will used to characterize the MD time series.

– **Lead time:** For a forecasting problem, the lead time specifies how far ahead the user wants to predict the future positions of atoms. Predicting far ahead (high lead) enables faster MD simulation, and at the same time, makes forecasting task more challenging.
– **History Size:** Next, we must decide how much historical data we wish to use to predict the future positions of atoms. This value is known as the History Size.
– **Prediction Window:** Prediction Window indicates the discrete time-window in the future used for creating the prediction outcome. For simplicity, in this paper, we always use a prediction window of 1 fs.
– **Prediction Error:** Error is defined as the Root Mean Square Deviation (RMSD (Eq.1)) between real and predicted structures at a given time point. During the learning stages, the error across individual interactions is denoted *loss*.

We present two new data sets to introduce subtleties in the equilibrium and nonequilibrium molecular dynamics from the perspective of time series forecasting. An analysis of these data sets will bring to light how effects of the history (or correlation) in the time series data can be described at different lead times and prediction windows to model a real-time dynamically evolving MD time series. The training objective here is to minimize the prediction error for a sufficiently large batch of training instances over a historical time span.

## 4 The New Data set

We introduce two data sets from two distinct kinds of MD simulation systems. Illustrated in Fig. 2, the first data set is an equilibrium simulation of the enzyme adenosine kinase (ADK). The second one is a steered molecular dynamics (SMD) or non-equilibrium simulation of the 100-alanine polypeptide helix (Fig. 3). In SMD, an external force is applied to the system along a chosen direction. We applied a force of 1 nanoNewton along one end of the 100-alanine helix, unfolding the protein[25].

**Fig. 2:**
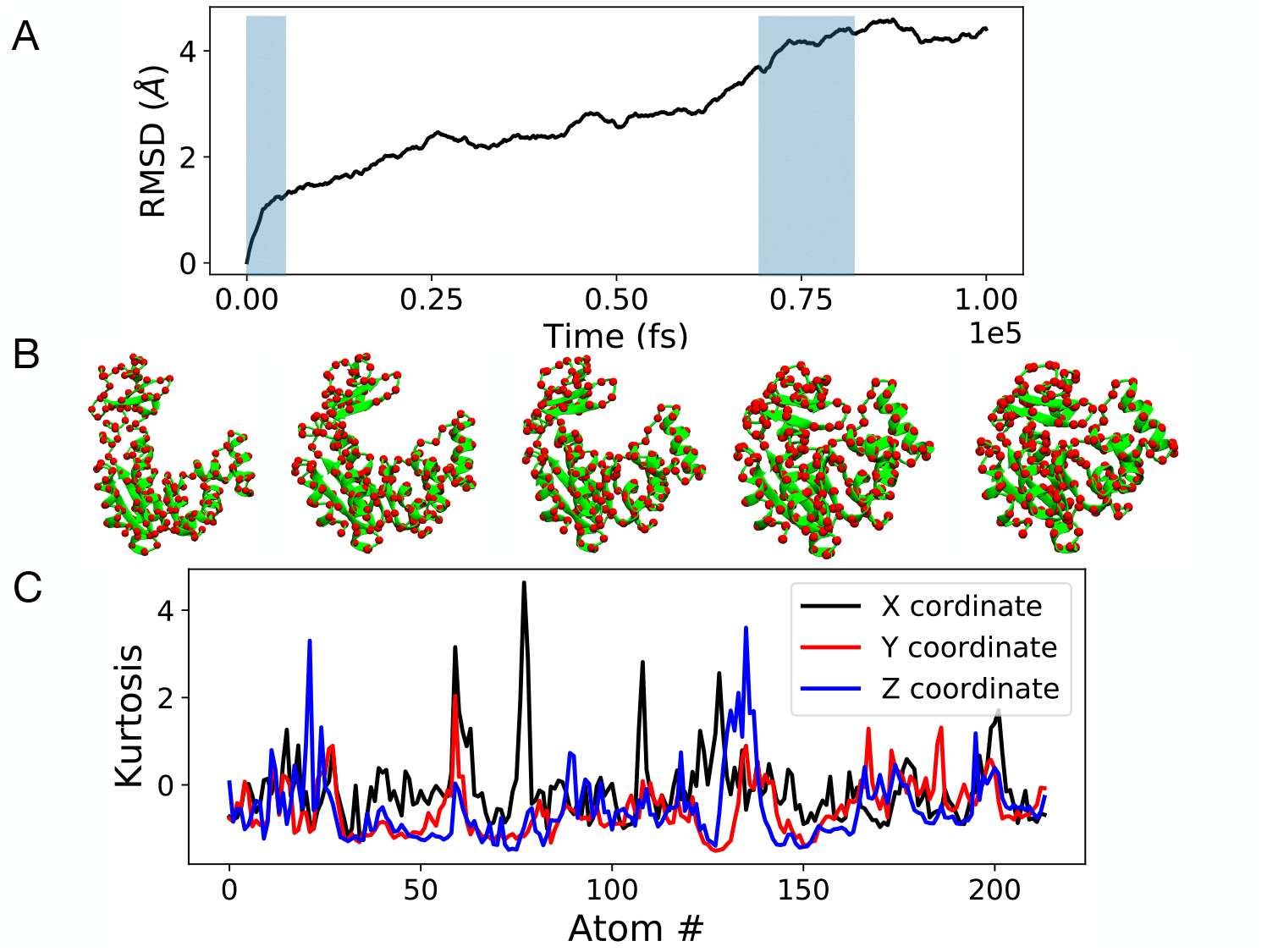
(A) RMSD of each time point of ADK equilbrium MD simulation with respect to the first time point, showing how the data varies over time (fs = femtoseconds). Regions of fast evolution are highlighted in blue. (B) Snapshots of the conformation of ADK at different time points of the trajectory (green: high dimension, red: reduced dimension) visualized in 3D and rendered in 2D using the molecular visualization software VMD[12]. (C) Deviation from Gaussian behavior (quantified by kurtosis where a higher value denotes larger deviation) of the distribution of X, Y, and Z positions of each of the 214 particles (shown in red in B).

**Fig. 3:**
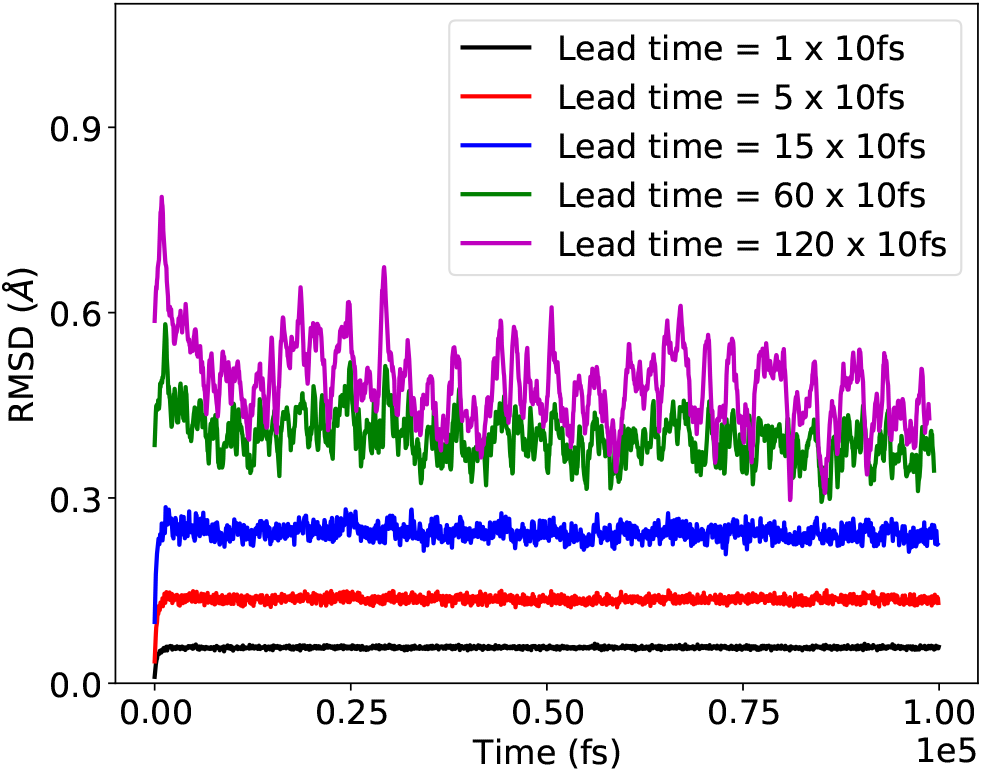
Error of forecasting from static model, at different lead times, for the ADK equilibrium MD simulation. Higher lead time makes the problem harder, thereby increasing the error.

We have generated high-as well as low-dimensional data for both the systems. In high-dimension, the position of every atom is explicitly defined, resulting in 3324 × 3 for ADK and 1003 × 3 dimensions for 100-alanine. For the low-dimensional data, positions of only the carbon atoms of each protein are defined, reducing the dimensionality of the problem to 214 × 3 and 100 × 3 respectively. The data is in X,Y,Z format presenting the Cartesean coordinates of the atoms for every time point along the time series. A total of 10^4^ time points is considered for the ADK example distributed evenly across 10^5^ fs (saved in steps of 10 fs - Figs. 2 and 3), and similarly 2000 data points were generated for the 100-alanine example across 10^7^ fs (saved in steps of 5000 fs - Figs. 4 and 5). The equilibrium time series was simulated employing OpenMM [6], while the non-equilibrium data set was constructed using our NAMD molecular dynamics simulation software packages [19].

**Fig. 4:**
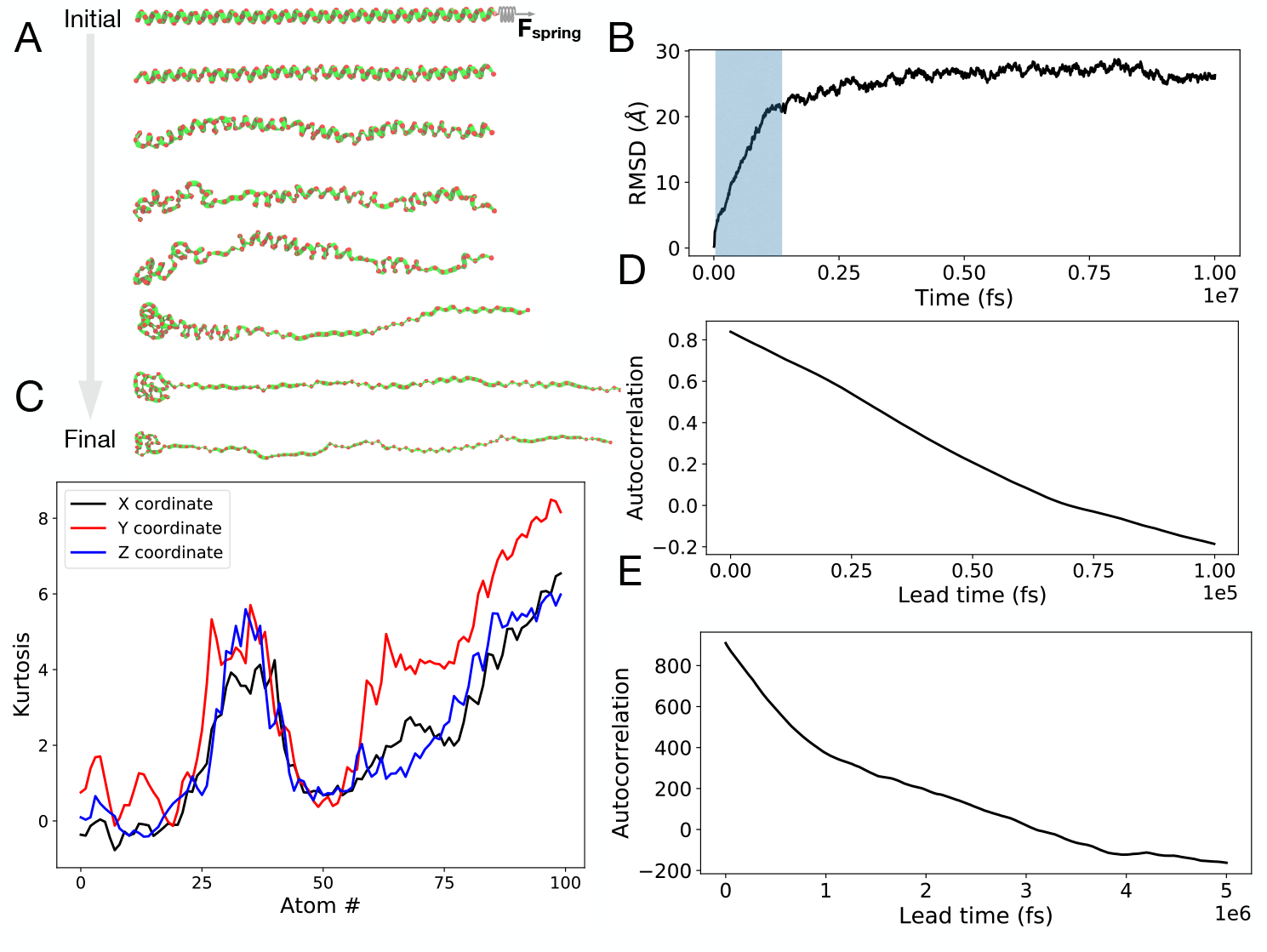
(A) Snapshots of structures from SMD trajectory at different time points (green: high, red: reduced dimension). (B) Deviation from Gaussian behavior (quantified by kurtosis, where a higher value denotes larger deviation) of the positions of the 214 particles (shown in red in A). (C) RMSD of 100-alanine with respect to the first time point, showing how the data varies over time. Regions of fast evolution are highlighted in blue. (D) Autocorrelation function of the radius of gyration (which reports on the shape-changes of the molecule) of ADK during the course of the simulation trajectory. Equilibrium MD simulation decorrelates in 10^5^ fs. (E) Autocorrelation function of the end-to-end distance (which reports on the extent of stretching) of 100-alanine during the course of the simulation trajectory. The non-equilibrium simulation takes 2-orders of magnitude more time to decorrelate.

**Fig. 5:**
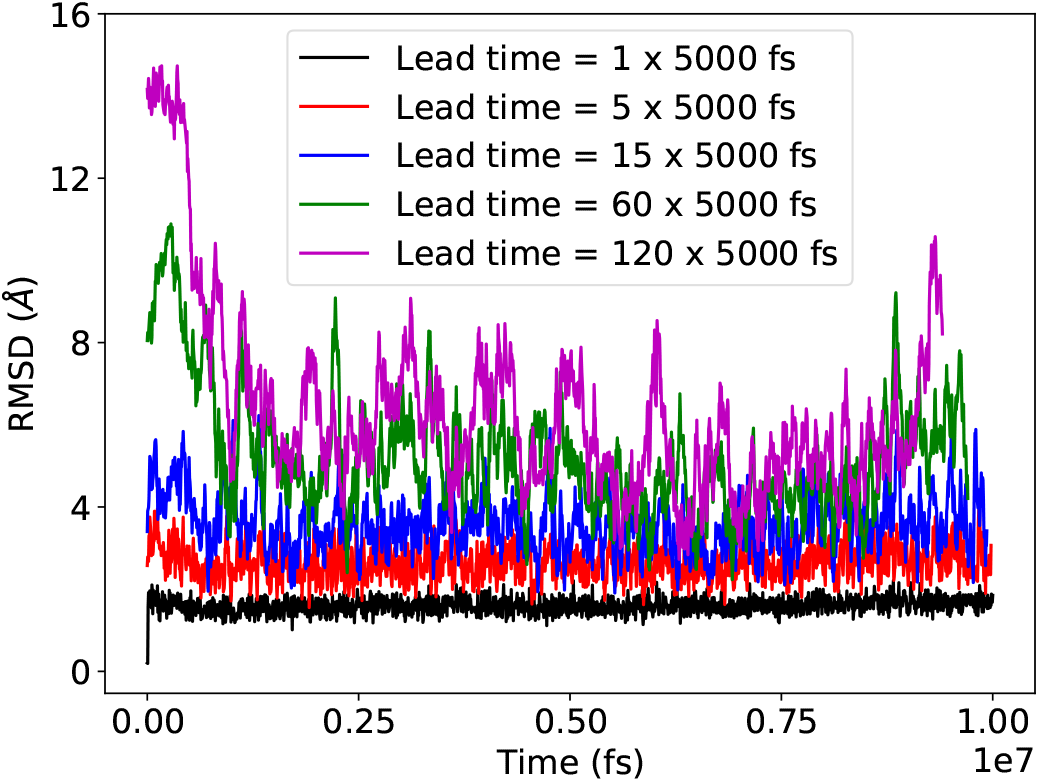
Error of forecasting from static model, at different lead times, for the 100-alanine SMD. Higher times make the problem harder.

In both the data sets, shape transformation of a 3-(3D) many-body system is recorded over time. For ADK, a transition from an open to a closed 3D-shape is observed due to concomitant rearrangements of 214 particles (Fig. 2B), while in 100-alanine, a more non-linear helix-to-coil transition is probed by tracking the changes in position of 100 particles (Fig. 4A). Beyond such high dimensionality of the data sets, the uniqueness of the equilibrium MD time series is in its dynamical evolution – the kinetic behavior stochastically switches between fast and slowly evolving regimes. Using RMSD values of all the the particle positions with respect to the very first, *t* = 0 position, we showcase these sudden changes in single-particle as well as collective dynamics in Fig. 2A.

For the non-equilibrium time series data of 100-alanine, the movements occur in the presence of an external force. These simulations produce less noisy data than the equilibrium MD of ADK Fig. 4B *vs.* 2A).

However, given that the shape changes are highly directed, we find that there are multiple classes of single-particle dynamics hidden under a collective behavior. Unlike the equilibrium MD simulations, where the positions of all the particles are Gaussian-distributed about a mean, at least two different classes of particle distributions is observed in the non-equilibrium time series (Fig. 4C *vs.* 2C). The distribution of the significant majority of atoms is nonGaussian, reflecting of the positional biases from high external forces to which they are subjected.

### 4.1 Equilibrium MD simulations of adenosine kinase (ADK)

During protein structure determination experiments, the atomic positions of a target protein are assigned by averaging the observed electron densities[9]. While this assignment offers a good starting model, the derived protein structure is typically in a non-biological (or non-native) state, and therefore severely limits biological application. Such artifacts can be resolved by bringing the starting model into thermal equilibrium at room temperature. Once in equilibrium, the protein adopts its native structure (3D shape) and dynamics. By numerically integrating Eq. 2, equilibrium MD simulations monitor the real-time evolution of native proteins.

The challenges involved in modeling of an equilibrium MD data can be presented employing the lead times of the associated time series. The hardness of the time series data is quantified by tracking how the RMSD values between the data points change at different lead times, namely at leads of 10,50..1200 fs (Fig. 3). The change in RMSD at different lead times also serve as a direct probe for the correlation in the data. If the lead time is short (10 or 50 fs) then it is simple to computationally probe the 0.1-0.2 Å scale changes in molecular position (Fig. 3, black and red traces) by analyzing the associated short-time correlations (Fig. 4D). In contrast, if the lead time is too long (600 and 1200 fs), then key short-time correlations within the data are missed. Thus, the associated small 3D shape changes may not be accurately learnt at this scale. One advantage of this data set is that all the particles are “well-behaved” and their dynamics is Gaussian distributed (Fig. 2C). Thus, an optimal lead time is desired which is sufficiently large (far into the future) to be interesting from a biological standpoint, and at the same time, can be used to train a machine learning model aimed at replacing computationally expensive MD.

#### Data preparation

A starting 3D protein model of ADK was generated using an x-ray diffraction crystal structure obtained from the PDB[2]. The atomic coordinates of ADK are encoded in the traditional PDB format presenting the X, Y, Z positions. X-ray is unable to resolve hydrogen atom positions. Thus, the position of hydrogen atoms were inferred using the run ADK.py script located in the *Equilibrium MD simulation* of the GitHub for this project [11]. Thus, a complete initial model was determined.

The goal of equilibrium MD is to recreate the native dynamics of a protein of interest. Therefore, the forces acting on each atom of the protein is defined using a potential energy function or force field.

The Amber force field, FF14SBonlysc, was used for the ADK simulation [6]. An implicit water model, GB-Neck2, was chosen to capture the equilibrium ADK environment; it is computationally efficient and enhances conformational sampling through decreased friction (*γ* in Eq. 2)[6]. After force field and water model selection, the energy of the protein model is minimized. The energy minimization corrects atoms that are in erroneously close contact due to artifacts from structural determination. If uncorrected, the simulation can produce unrealistic forces that cause the simulation to become unstable. Once minimized using conjugate gradients, the all-atom model is ready for production simulation.

The ADK simulation was performed for 10^5^ timesteps with a periodic update frequency of 1 fs, and atomic models were saved every 10 fs. This results in a 0.1 nanosecond (10^5^ steps × 1 fs/step) simulation of the ADK protein, providing in time series of 10^4^ data points. The simulation of ADK was performed using the openMM python library[6]. Five copies simulations were performed at a temperature of 310K. Collective dynamics of ADK was monitored by computing its RMSD relative to the *t* = 0 time point (Fig. 2A). A plateau in this profile suggests that equilibrium is attained at 0.8 × 10^5^ fs. The trajectory data, containing 10^4^ time points or snapshots, was initially stored in single precision binary FORTRAN files known as DCD files. The positional coordinates (X,Y,Z) of all atoms in each snapshot were extracted from the DCD file resulting in a rank-3 tensor which was (3324 × 3 × 10^4^) for the high dimensional space and (214 × 3 × 10^4^) for the low dimensional data. The entire simulation can be reproduced with a single OpenMM python script located in the *Equilibrium MD simulation* on GitHub [11].

### 4.2 Non-equilibrium MD simulations of 100-Alanine

Life as we know, exists out of equilibrium. Traditionally, experiments focusing on the nonequilibrium behavior of proteins were performed by either adding heat or inducing chemical perturbations. Another factor that can bring proteins out of equilibrium is mechanical stress (e.g stretching of the molecules). Such stretching arises naturally in proteins located in the muscle tissue. The response of these proteins to mechanical stress can be studied by investigating the individual domain’s response to stretching within Atomic Force Microscopy or AFM experiments[16]. This molecular events are analogous to the process of pulling a rubber-band by holding one end fixed in our hand (Fig. 4A).

Now, we employ non-equilibrium MD simulations for computationally recreating the AFM experiments. In particular, Steered MD or SMD is used to generate a relevant and challenging data set for learning algorithms to be trained and validated. It is notable that events from such non-equilibrium pulling experiments or their equivalent SMD simulations, have never been used within RNN, in particular LSTM frame-work for time series forecasting.

The challenge in SMD is commensurate to that of equilibrium MD in that, an optimal lead time should be derived respecting the correlation limits of the data. However, subtleties are twofold: first, for the same lead time steps the RMSD error bars in SMD are much higher (Fig. 5), consistent with more prominent 3D shape changes that those observed for equilibrium MD simulations of ADK (Fig. 4A *vs.* 2B). Yet, the longer the correlation times (Fig. 4E) indicate smoother shifts within the time series. Second, there are multiple classes of atoms with different dynamics distribution (Fig. 4C).

#### Data preparation

The 100-alanine helix was prepared using the Avagadro software on a single CPU. The external force acts on the C-terminus of the long helical protein, while the N-terminus region remains constrained. As the molecule is stretched, it undergoes a gradual conformational change, transitioning from an *α*-helix to a random coil (Fig. 4A). Typically, there are two variants of SMD, constant force and constant velocity pulling. The equation for the external pulling force (*F_spring_*) acting on the atom in the C-terminal region of the protein is given by,

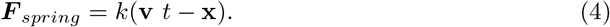

Here, **x** is the displacement of the atom in protein which is pulled from its original position, **v** is the prescribed pulling velocity, and k is the spring constant. In the presence of this external force, the equation of motion (Eq. 2) becomes

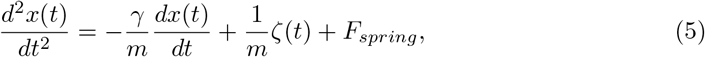

For our data set, we adopt the SMD with constant velocity (SMD-CV) protocol from our open-source NAMD tutorial[25]. The SMD-CV simulations are performed using the Langevin dynamics scheme of MD at constant temperature of 300 K in Generalized-Born implicit solvent with the CHARMM36m force field[19]. One end of the molecule (N-terminus) is constrained while the other end at the (C-terminus) is free to move along the z-axis with a constant speed of 0.2 Å/ps and force constant of 7 kcal/mol/Å^2^, exerting an overall force of 1 nanoNewton (Fig. 4A)[16].

A set of 5 copies of SMD is used to generate an ensemble of conformations when subject to SMD-CV pulling. All simulations are performed using the recent build of NAMD (version Nightly Build) with a time step of 1 fs, with dielectric constant of 20, and a user defined cut-off for Coulomb forces with a switching function starting at a distance of 10 Å which plateaus to zero at 12 Å. A simulation time of 10^7^ fs is required for extension of the helix to random coil. Here, we save the trajectory every 10 fs, mainly to generate a large data set of 10^6^ points to train an LSTM model in Sect. 5. The data presented in Figs. 4 and 5 are saved at even longer time intervals, namely 5000 fs, to reduce the number of time points to 2000 for computing lead times and correlations. The full data set of (1003 or 100 × 3 × 10^6^) points, which is used in the LSTMs below is accessible through the google drive link provided on GitHub [11]. A Tcl script *smd.constvel.namd* is used to implement the outlined simulation protocol. The script includes all the standard NAMD parameters, which are outlined above to perform SMD. This script together with all other input files are available freely through GitHub [11] and the NAMD website[25] to reproduce our data set for non-equilibrium MD simulations.

### 4.3 Utility and Predicted Impact

Our MD data is documented in tutorial files, scripts, and an openly accessible GitHub page [11] so any user with access to a single CPU or GPU node will be able to reproduce the results. The full time series can be loaded, visualized in 3D and analyzed for RMSD using the molecular visualization tool VMD (Figs. 2 and 4).

The presented data set exemplifies arguably a first attempt at capturing the entire range of time series variations typical of a biomolecule. We describe two broad classes of data with distinct correlation timescales. More importantly, the data clearly shows how external physical forces can alter time series correlations and provides an avenue to experiment with machine learning models for probing such external factors. Accordingly, a data scientist can chose a suite of different learning algorithms to model these fast evolving high dimensional MD trajectory data.

The equilibrium data at a single-particle level appears to be well behaved with relatively uniform Kurtosis values (Fig. 2C), but offers difficulties in training of the rapid variability in RMSDs (Fig. 2A–multiple shaded regions).

In contrast, the non-equilibrium data shows non-Gaussian statistics at a single-particle level (Fig. 4C) eliciting complexity at a single-particle level, but manifest smooth changes in the time series when treated together (Fig. 4B).

A key question these data sets pose is whether a common learning algorithm can ever be introduced to work with all the limits of biomolecular dynamics. A second question the data sets raise pertains to identification of limits that are easier to model using popular sequence modeling techniques like RNNs with LSTMs or GRUs cells either in isolation or in concert. Finally, will the learning algorithms scale if the dimensions of the data sets increase from the hundred-to-thousand variables, chosen here for simplicity, to the more realistic million-to-billion dimensional spaces. These three questions also offer the opportunity to think about the use of the existing petascale or the upcoming exascale resources for handing the convoluted biomolecular problems with data science methodologies. Put together, these data sets places an machine learning expert in a position to address one of the central questions at the interface of life sciences and computer sciences, namely to what extent can numerical simulation schemes be by-passed using the machine learning tools. The community of computational biophysics with nearly 20,000 NAMD users and a 3-4 fold large cadre of researchers applying MD will immediately benefit from answering this question. The findings from this data set are further generalizable to any domain with quantitative data on high-dimensional rapidly fluctuating time series.

## 5 An Exploratory Study

Due to the recent success of Recurrent Neural Networks (RNN) for modeling time series data [4], we conducted an exploratory study with RNNs to model the two new dynamically evolving MD trajectory data. We used Long-Short Term Memory (LSTM) cells in the hidden layers and trained RNNs on both equilibrium and non-equilibrium MD simulations to decipher which data-set is more amenable to learning. More specifically, we conducted a series of experiments to produce baseline accuracy numbers for LSTMs as well as to tune the different hyperparameters associated with the same. Below we present a brief summary of the experiments that were conducted and report our findings to facilitate in-depth future research in this direction.

### 5.1 Setting a Baseline: The Static Model

As a starting point, we set the *Static Model* as our baseline where we assume that the position of an atom at a future timestamp *X_t_*_+*lead*_ does not change relative to its last known position, i.e. *X_t_*, where, *t* is the current timestamp. The assumption is incorrect, but still helps us set a realistic baseline for evaluating the performance of advanced machine learning techniques like LSTMs. Figures 6A,B (ADK) and 8A,B (SMD) show the RMSD distributions of static model for lead time steps 15 and 120, respectively.

**Fig. 6:**
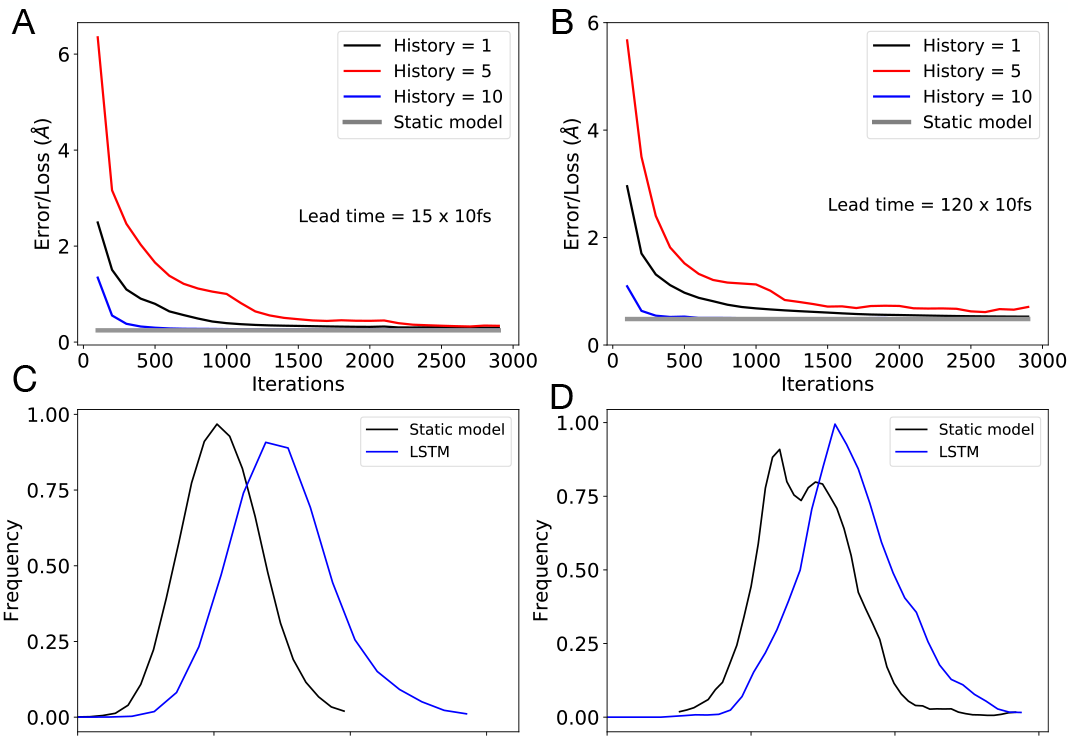
Training loss for ADK equilibrium MD simulation, as a function of history for (A) lead time step = 15 and (B) lead time step = 120. Black, red and blue lines represent, respectively, history of 1, 5 and 10 time points. Distribution of RMSD for LSTM prediction compared to that of static model prediction for (C) lead time step = 15 and (D) lead time step = 120. For ADK equilibrium MD simulation, LSTM performs poorer than static model.

### 5.2 Training LSTM

For starters, we trained a RNN with 32 LSTM units in the hidden layer, a learning rate of 0.01, history size 5 and varying lead time steps of {1, 5, 15, 60, 120}. The output layer used linear activation and Mean Squared Loss was used as the training loss function. Below we report some of our key observations from the experiments.

#### Curse of Dimensionality and Kirchhoff decomposition

We found that learning by treating the entire protein structure at a given timestamp as a single training instance is very challenging due to the high dimensionality of the problem, generating higher errors than the static model. To deal with this issue, we assumed that the position of each atom within the protein structure is independent of one another and can be modeled as separate one dimensional time-series. This so-called Kirchhoff decomposition scheme boosted the performance of LSTM significantly.

##### ADK Vs 100-alanine

We report RMSD of each simulated system, i.e., ADK and SMD (Figs. 2A and 4B). We found that RMSD of SMD simulation of 100-alanine is one order of magnitude higher than that of the equilibrium MD simulation of ADK. This is due to the non-equilibrium nature of the former, where an external force is used to pull the system. This difference is also reflected in the static model error at varying lead time steps (Figs. 3 and 5).

##### Effect of Lead time

Increasing lead time makes time series forecasting harder, which we expected would justify the use of complex sequence modeling techniques like LSTM. In other words, we hypothesized that an increase in lead time will cause the LSTM error to increase less than the static model error. We found this to be true for the 100-alanine SMD simulation. With lead time steps of 1 and 5, LSTM loss was higher than the static model error. However, with lead time step of 15, LSTM performed better than the static model, and improvement from static model increased further at even higher lead time steps (60 and 120). Due to lack of space, we only present the results for lead time 15 an 120 (Fig. 8 A,B).

**Fig. 7:**
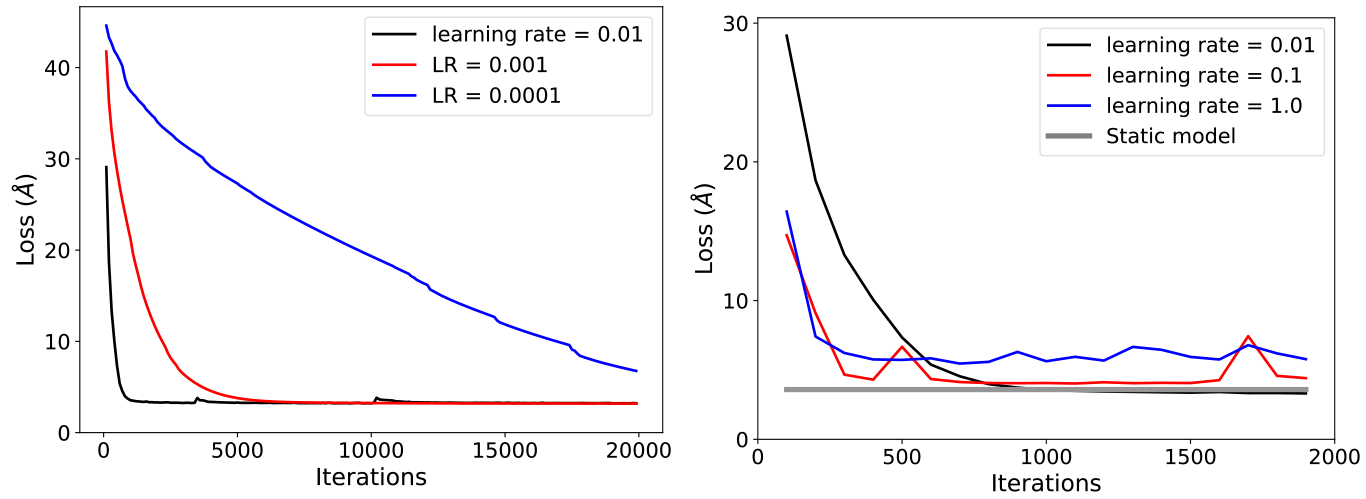
Effect of learning rate on LSTM training on SMD trajectories.

**Fig. 8:**
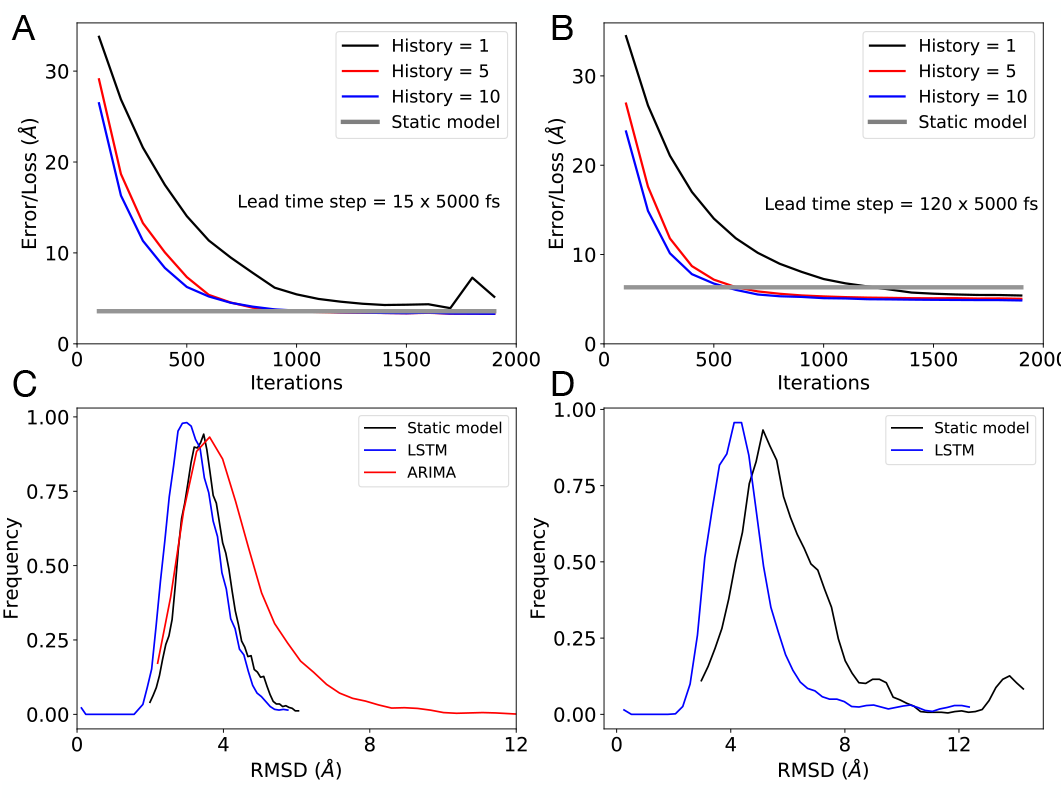
Training loss for 100-alanine SMD simulation, as a function of history for (A) lead time step = 15 and (B) lead time step = 120. Black, red and blue lines represent, respectively, history of 1, 5 and 10 time points. Distribution of RMSD for LSTM prediction compared to that of static model prediction for (C) lead time step = 15 and (D) lead time step = 120. For 100-alanine SMD simulation, LSTM performs better than static model, and performance improves at higher lead time. For comparison, we also report the result of ARIMA (red) with a lead time step of 15.

In contrast, we have not been able to achieve lower LSTM losses compared to the static model loss for the equilibrium MD simulation of ADK, for the lead time steps 1 through 120. Equilibrium MD simulation of ADK decorrelates much faster than SMD, in the picosecond regime (Fig. 2A). This yields an interesting as well as surprising result that equilibrium MD trajectories were more difficult to model than the non-equilibrium MD trajectories, which is indeed counter intuitive.

##### Effect of History

For this set of experiments, we hypothesized that an increase in history size will reduce the LSTM training error as we are using more information from the past. Indeed, the results confirm our hypothesis (Figs. 6AB and 8A,B). More Specifically, we varied history size among {1, 5, 10} and found that increasing the history actually reduces the LSTM training erros for both ADK and 100-alanine trajectories.

##### Effect of learning rate

We trained the LSTM network separately while varying learning rate among {1.0, 0.1, 0.01, 0.001, 0.0001}. We found that rates of 1.0, 0.1 were unstable, while 0.001, 0.0001 were too slow to converge for SMD simulations (Figs. 7) [Results for MD simulations were similar, and are provided in GitHub[11]]. Thus, we recommend 0.01 as the learning rate.

##### Summary of Hyper-parameter Tuning Study

Based on our exploratory study, we recommend the following set of empirical values for each hyper-parameter as shown in Table 1.

**Table 1.**
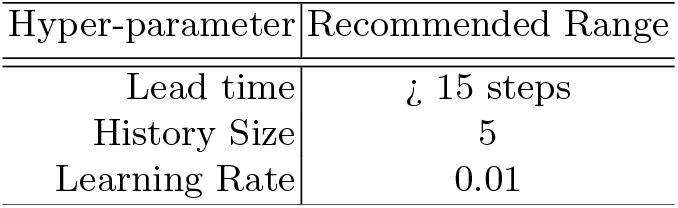

## 6 Future Directions

Table 1: Recommended range of hyper-parameters. 32 hidden units were used.

In regards to the future directions of methods that can be done in the data set, there are still more ways to improve training on LSTMs. One possible improvement is through more stacked LSTMs. This would be able to learn more nonlinear dynamical relationships between the points.

Other than LSTMs, we can also borrow from deep learning in natural language processing by utilizing attention models, which have recently been getting state of the art results, without of the use of a recurrent hidden layer [4].

Other considerations for future direction is the ability to reformulate the 3D structural input of the data as a 3D point cloud. There have been recent deep learning architectures used in 3D point cloud segmentation and classification such as VoxelNet and PointNet [3]. Both architectures leverage the underlying 3D relationship between points and objects in 3D space for the supervising task. With VoxelNet, the data is voxelized into fixed voxels in which a 3D convolutional neural network is used. However, with architectures like PointNet, the input can be variable. In this case, future directions can be the addition of a data set in which the number of atoms per dynamical system and be varied.

With architectures that deal with data in the 3D space, there is the consideration of new loss functions. Here, we utilized MSE loss in optimizing our LSTM. Loss functions such as Earth Movers Distance (EMD) and Chamfer Loss are two most notable losses used for 3D point generation [8]. Moreover, EMD can be extended for graphs, which can be useful for not only learning the 3D geometrical relationships, but the graph relationships between atoms.

The external information sought in the current data sets from AFM or force measurements to improve temporal correlation can also be derived from other experimental modalities such as X-ray crystallography [18] or cryo-electron microscopy [24]. Finally, recovery of the all-atom description from an LSTM-predicted reduced space of only heavy atoms opens the door to inverse-Boltzmann approaches for reverse coarse-graining [9].

## 7 Conclusion

In the present study, we report two new data sets for describing equilibrium and non-equilibrium protein dynamics produced by physics-based simulations. These data sets fill a much needed knowledge gap in the protein-learning field, providing a synergistic augmentation to the popular existing data sets used for learning molecular structure [2]. Protein dynamics was represented as a time-series data and was modeled through a recurrent neural network with LSTM cells in the hidden layer. We found that the learning of both data sets was improved when using a Kirchhoff decomposition on models with a constant number of hidden layers. The ability to forecast future structure was shown to be dependent on the correlation among the recent past structures. Specifically, dynamics within the non-equilibrium molecular dynamic simulations were highly correlated, and thus protein dynamics were effectively learned. Conversely, the movements of a protein at thermal equilibrium were poorly correlated, making accurate forecasting more difficult. Increasing history size improved the prediction accuracy for both data-sets and LSTM outperformed the static baseline while forecasting at higher lead times. Overall, LSTMs provide an exciting tool to model non-equilibrium protein dynamics. Virtually all biologically relevant actions occur at non-equilibrium, therefore these results indicate an exciting advance with far-reaching implications.

## Notes

### Competing Interest Statement

The authors have declared no competing interest.

